# Sexually dimorphic differentiation of a *C. elegans* hub neuron is cell-autonomously controlled by a conserved transcription factor

**DOI:** 10.1101/081083

**Authors:** Esther Serrano-Saiz, Meital Oren-Suissa, Emily A. Bayer, Oliver Hobert

## Abstract

Functional and anatomical sexual dimorphisms in the brain are either the result of cells that are generated only in one sex, or a manifestation of sex-specific differentiation of neurons present in both sexes. The PHC neurons of the nematode *C. elegans* differentiate in a strikingly sex-specific manner. While in hermaphrodites the PHC neurons display a canonical pattern of synaptic connectivity similar to that of other sensory neurons, PHC differentiates into a densely connected hub sensory/interneuron in males, integrating a large number of male-specific synaptic inputs and conveying them to both male-specific and sex-shared circuitry. We describe that the differentiation into such a hub neuron involves the sex-specific scaling of several components of the synaptic vesicle machinery, including the vesicular glutamate transporter *eat-4/VGLUT,* induction of neuropeptide expression, changes in axonal projection morphology and a switch in neuronal function. We demonstrate that these molecular and anatomical remodeling events are controlled cell-autonomously by the phylogenetically conserved Doublesex homolog *dmd-3,* which is both required and sufficient for sex-specific PHC differentiation. Cellular specificity of *dmd-3* action is ensured by its collaboration with non-sex specific terminal selector-type transcription factors whereas sex-specificity of *dmd-3* action is ensured by the hermaphrodite-specific master regulator of hermaphroditic cell identity, the Gli-like transcription factor *tra-1*, which transcriptionally represses *dmd-3* in hermaphrodite PHC. Taken together, our studies provide mechanistic insights into how neurons are specified in a sexually dimorphic manner.

## INTRODUCTION

Male and female brains display sexual dimorphisms on the level of function, anatomy and gene expression. In principle, such dimorphisms can be a reflection of two distinct scenarios; one in which dimorphisms are the result of the presence (or expansion of numbers) of neurons in a specific brain area specifically in one sex but not the other, while in the other scenario neurons that are present in both sexes (“sex-shared neurons”) acquire sex-specific features [1]. These scenarios are hard to untangle in vertebrate nervous systems, since their cellular complexity confounds the visualization and systematic comparison of individual neuron types between the two sexes. In contrast, clear evidence exists for both scenarios in the *Drosophila* nervous system, that is, both sex-specific neurons, as well as sex-specific anatomical features of sex-shared neurons have been unambiguously mapped [1-3]. However, dimorphic features of shared neurons have so far been only studied on a relatively coarse level and, aside from well-characterized sex-specific regulatory factors (Fruitless and Doublesex; [4]), there are relatively few molecular and functional features that have been mapped onto sex-shared, but sexually dimorphic neurons.

In the nematode *C. elegans,* sexually dimorphic features have been analyzed with unprecedented anatomical and molecular resolution [5-8]. Like in other organisms, the *C. elegans* nervous system contains sex-specific neurons found exclusively in either the male or the hermaphrodite (a somatic female). Moreover, there are neurons that are present in both sexes that share an identical lineage history, position, molecular features and overall appearance but that display sexually dimorphic synaptic connectivity patterns [6]. In one set of striking examples, the sex-shared phasmid neurons PHA and PHB connect in a sexually dimorphic manner to distinct downstream interneurons which are also sex-shared. We have recently shown that dimorphic synaptic connections among shared neurons mostly arise through synaptic pruning processes [9].

Apart from sexual dimorphisms of synaptic connectivity between sex-shared neurons, there are also sexually dimorphic synaptic connections between sex-shared neurons and sex-specific neurons [6]. Relatively few sex-shared neurons receive synapses from sex-specific neurons and these neurons can be considered “hub neurons” since they connect sex-specific network modules to the shared and non-sex specific “core nervous system” [6]. The most remarkable examples of such hub neurons are the PHC sensory neuron, and the DD6 and PDB motorneurons. All three neurons are sex-shared and receive a tremendous amount of additional synapses from male-specific neurons [6]. PHC stands out among those, because unlike DD6 and PDB, it is not a motor neuron that directly connects male-specific sensory inputs into muscle. As schematically diagrammed in Fig.1A, PHC rather connects in a sexually dimorphic manner to a diverse set of inter- and motorneurons and of all neurons in the animal, it is the neuron that receives the most male-specific synaptic inputs from a variety of different sensory neurons. This is in notable contrast to the synaptic connectivity of PHC in hermaphrodites, where PHC appears a canonical sensory neuron receiving little synaptic inputs and making modest outputs on a number of interneurons (Fig.1A)[5]. Taken together, PHC undergoes a sexual differentiation process in males that transform this neuron from a conventional sensory neuron to a hub neuron with sensory and interneuron properties.

**Fig. 1:**
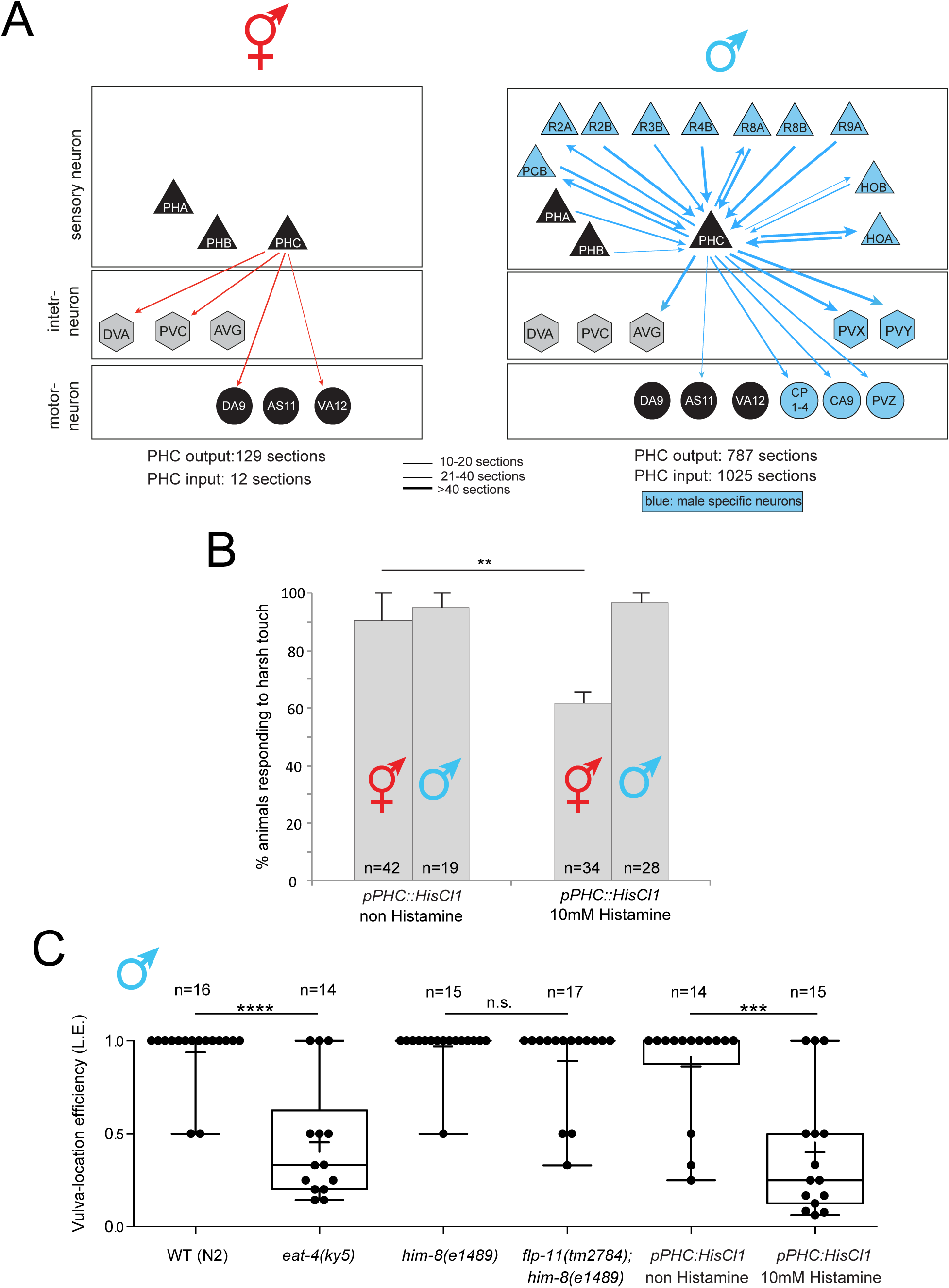
Dimorphic synaptic connectivity and function of the PHC neuron. **A:** Schematic synaptic connectivity. Edge weights were collected from www.wormwiring.org [6]. Only connections that directly involve PHC are shown. **B:** PHC is required in hermaphrodites for response to harsh (picking-force) touch to the tail. Bar graph indicates the mean percentage of animals responding to harsh touch, with error bars indicating the minimum and maximum from two experimental replicates. Significance was calculated using Fisher’s exact test. ** p<0.01, n.s. p>0.05. “n” in each column indicates the total number of animals assayed across two experimental replicates. **C:** PHC is required for a specific step of the male mating behavior, vulva search behavior. Mutant and Histamine-silenced animals tested for the male’s vulva location efficiency. Box plot representation of the data is shown, with whiskers pointing from min to max. Median and mean values are indicated by horizontal line and “+”, respectively. Statistics was calculated using Kruskal-Wallis test. \*\*\*\**P* < 0.0001, \*\*\**P* < 0.0005. “n” above graphs indicates the total number of animals assayed.

In this manuscript we assign sex-specific functions to PHC and describe a number of distinct molecular features of the PHC neurons, which parallel these strikingly distinct anatomical features. These include the scaling of expression of synaptic vesicle components, a phenomenon not previously described. We show that anatomical and molecular dimorphisms are superimposed during sexual maturation of the male onto an initially non-dimorphic morphology and gene expression program. Furthermore, we show that this remodeling is controlled in a cell-autonomous manner by two phylogenetically conserved transcription factors, TRA-1/Gli, a global regulator of sexual identity in *C. elegans* and DMD-3, a member of a phylogenetically conserved family of DM domain transcription factors, defined by Doublesex in Drosophila and DMRT genes in vertebrates. This family of transcription factors represents the only unifying theme of the otherwise very divergent regulatory mechanisms of sex determination and sexual differentiation throughout the animal kingdom [10].

## RESULTS

### Sex-specific differences in PHC neuron function

We first set out to define a behavioral consequence of the male-specific rewiring of the PHC neuron. We generated transgenic animals in which the PHC neurons can be silenced in an inducible manner with a histamine-gated chloride channel [11]. Based on its direct innervation of command interneurons, a unique feature of several, previously described nociceptive neurons like ASH, ADL or PHB [5, 12], we tested the response of these transgenic animals to nociceptive stimuli and found that silencing of PHC results in defects in the response to harsh touch applied to the tail of the hermaphrodite (Fig.1B). In contrast, silencing of PHC in males has no effect on the tail harsh touch response. This functional difference suggests that another neuron is now responsible for this response in the male tail and it is consistent with both dramatic changes in dendrite morphology of male PHC (described below) and the absence of PHC innervation of command interneurons in males (Fig.1A).

We found that male PHC becomes repurposed for male mating behavior. Specifically, males treated with histamine showed profound defects in vulva location behavior. Instead of stopping when the male tail has reached the vulva, PHC-silenced animals continue to search for the vulva (Fig.1C). This phenotypic defect is consistent with the PHC wiring pattern: two sensory neurons that innervate PHC, the PHB and HOA, as well as an interneuron (called AVG) innervated by all three neurons, PHC, PHB and HOA, were previously shown to be involved in responding to a vulval stop signal [9, 13]. Similar defects can be observed upon genetically disrupting the neurotransmitter system, glutamate, used by all these neurons (PHB, PHC, HOA) (Fig.1C).

### Sex-specific, cell-autonomous morphological differentiation of the PHC neurons

Having established a sex-specific function of PHC, we set out to define the sex-specific differentiation program of PHC in more detail. The electron micrographic analysis of two adult hermaphrodite tails and one adult male tail indicates that apart from their notably distinct synaptic connectivity patterns, the PHC neurons display different axon and dendrite lengths [5, 6, 14]. We used a reporter transgene, described in more detail below (*eat-4prom11Δ12*), to visualize the sexually dimorphic morphology of PHC in a larger cohort of animals, and we examined its previously undocumented dynamic development during larval stages (Fig.2A). We find that up until the L3 stage, before sexual maturation, PHC extends its dendrite into the tail tip of both sexes (Fig.2A, purple arrow). During retraction and remodeling of the male tail hypodermis, the PHC dendrite then retracts and adopts a meandering morphology (Fig.2A). The axons of the PHC neurons extend initially only into the pre-anal ganglion of both sexes. However, beginning at L4 stage, when male-specific neurons are generated and start to differentiate, the axon of male but not hermaphrodite PHC neurons extends significantly beyond the pre-anal ganglion into the ventral nerve cord (Fig.2A, B), to then gather synaptic inputs from male-specific neurons and innervate newly generated male-specific motor neurons as well as sex-shared neurons (Fig.1A)[6].

**Fig. 2:**
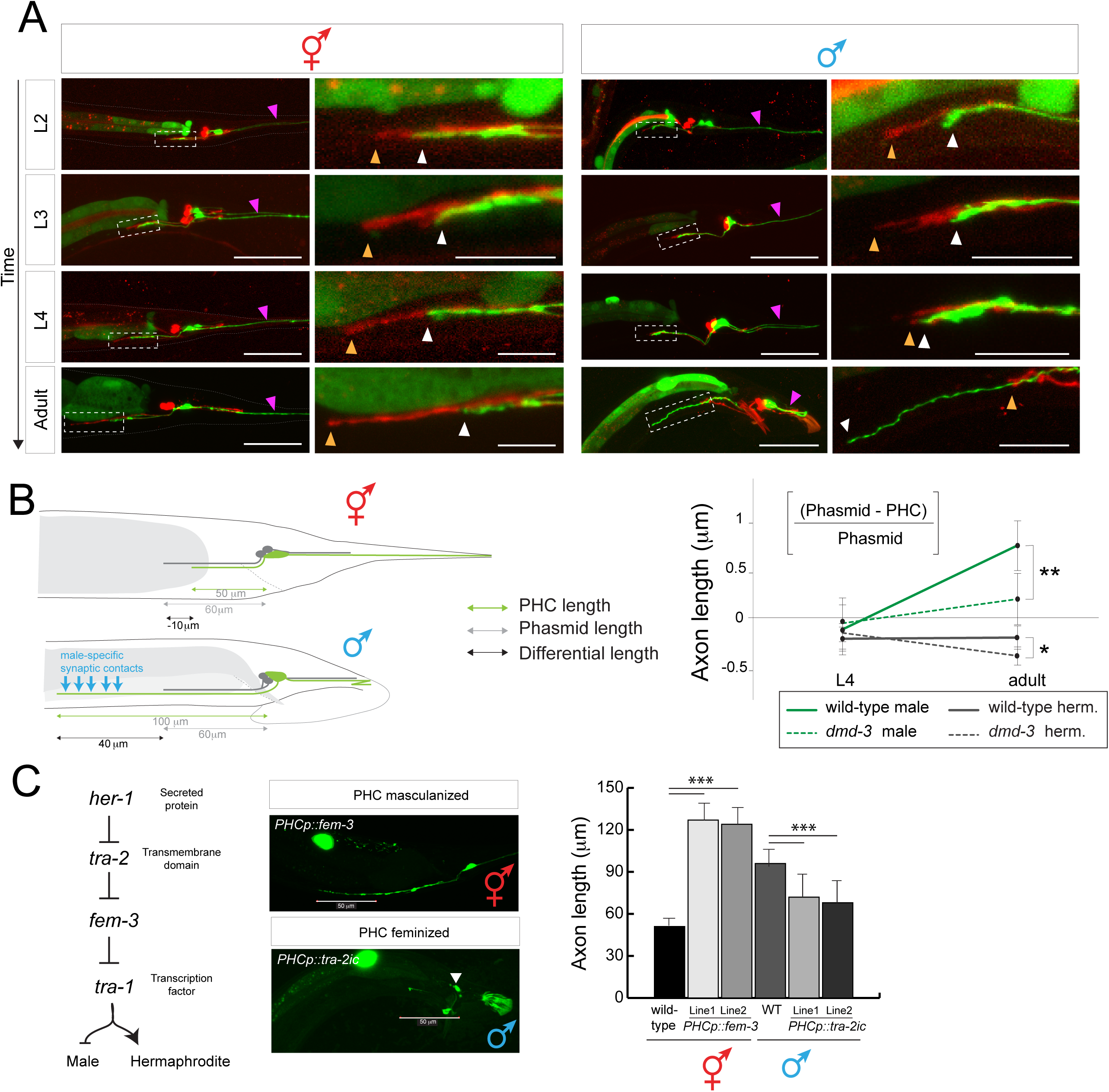
Sexually dimorphic extension of the PHC axon. **A:** PHC axon and dendrite morphology at different developmental stages. PHC was visualized with the transgene *otEx6776*, which expresses *gfp* under the control of the *eat-4prom11Δ12* driver, which expresses in PHC of both males and hermaphrodites (described in more detail in Fig.5). As landmark, the PHA and PHB neurons, which do not noticeably change morphology between the sexes, are filled with the red dye DiD (termination point of PHA/B marked with orange arrowhead, PHC with white arrowhead). PHC dendrites are marked with purple arrowhead. Scale bar: 50 µm. Dashed inbox indicates the area that is magnified in the left column. Scale bar in magnified panels: 10 µm. **B:** Summary of PHC axon extension. Male specific synaptic contacts (referring to both inputs and outputs) are schematically indicated with blue arrows. **C:** Male-specific axon extension is controlled cell-autonomously as determined by cell-specific sex change via manipulation of the sex-determination pathway. Transgenic array names: *otIs520* (*eat-4prom11::gfp*) was crossed with *PHCp::fem-3* (lines *otEx6879* and *otEx6980*) and *PHCp::tra-2ic (lines otEx6881 and oEx6882)* strains. PHCp (*eat-4prom11Δ11* driver). Significance was calculated using student t-test, \*\*\**P* < 0.0005, \*\**P* < 0.005 and \**P* < 0.01.

We asked whether this sex-specific extension of the PHC neurons depends on male-specific synaptic targets which may secrete attractive cue(s) or on any other male-specific structure or whether sex-specific PHC axon extension is controlled in a cell-autonomous manner. To this end, we altered the sex of the PHC neurons in a cell-autonomous manner through manipulation of the activity of the Gli-like transcription factor TRA-1, the master regulator of sexual differentiation [15]. TRA-1 is expressed in all hermaphroditic cells and required autonomously to promote hermaphroditic cellular identities and repress male identities [16]. In males, TRA-1 is downregulated via protein degradation (see schematic in Fig.2C)[17]. We feminized male PHC neurons through PHC-specific expression (*eat-4prom11Δ11* driver, from now on called *PHCp*) of the intracellular domain of TRA-2, *tra-2ic*, which constitutively signals to prevent TRA-1 downregulating by FEM-3 and other factors [18, 19]. We find that in otherwise male animals this cell-autonomous feminization results in a failure of the PHC axon to undergo its characteristic extension through the pre-anal ganglion (Fig.2C).

Conversely, masculinization of PHC via *fem-3*-mediated degradation of TRA-1 protein in otherwise hermaphroditic animals results in a male-like extension of the PHC axon (Fig.2C). These sex-reversal experiments confirm cell autonomy but also demonstrate that (1) the TRA-1 transcription factor represses a male morphological differentiation program in PHC and (2) that TRA-1 is not only required to repress male-specific features in hermaphroditic PHC, but is apparently also sufficient to repress male-specific features of PHC.

### Transcriptional scaling of glutamatergic neurotransmission in the male PHC neuron

We find that sex-specific morphological changes of the PHC neurons are paralleled by a striking change in a number of molecular features. Specifically, we noted that the expression of the vesicular glutamate transporter *eat-4/Vglut* fosmid is strongly upregulated in the PHC sensory neurons, as assessed with a fosmid reporter construct (Fig.3A). We quantified this upregulation through normalization of *eat-4/Vglut* expression relative to *eat-4/Vglut* expression in other neurons, and also relative to a ubiquitously expressed nuclear envelope protein-encoding gene, *mel-28* (Fig.3B). We independently validated this upregulation using quantitative single mRNA fluorescence in situ hybridization (smFISH)(Fig.3D). Both reporter gene and smFISH analysis suggest a threefold upregulation in *eat-4/Vglut* expression (Fig.3). Assessing the timing of the upregulation of *eat-4/Vglut* expression, we find it to begin at the late L4/young adult stage (**Fig.3C**).

**Fig. 3:**
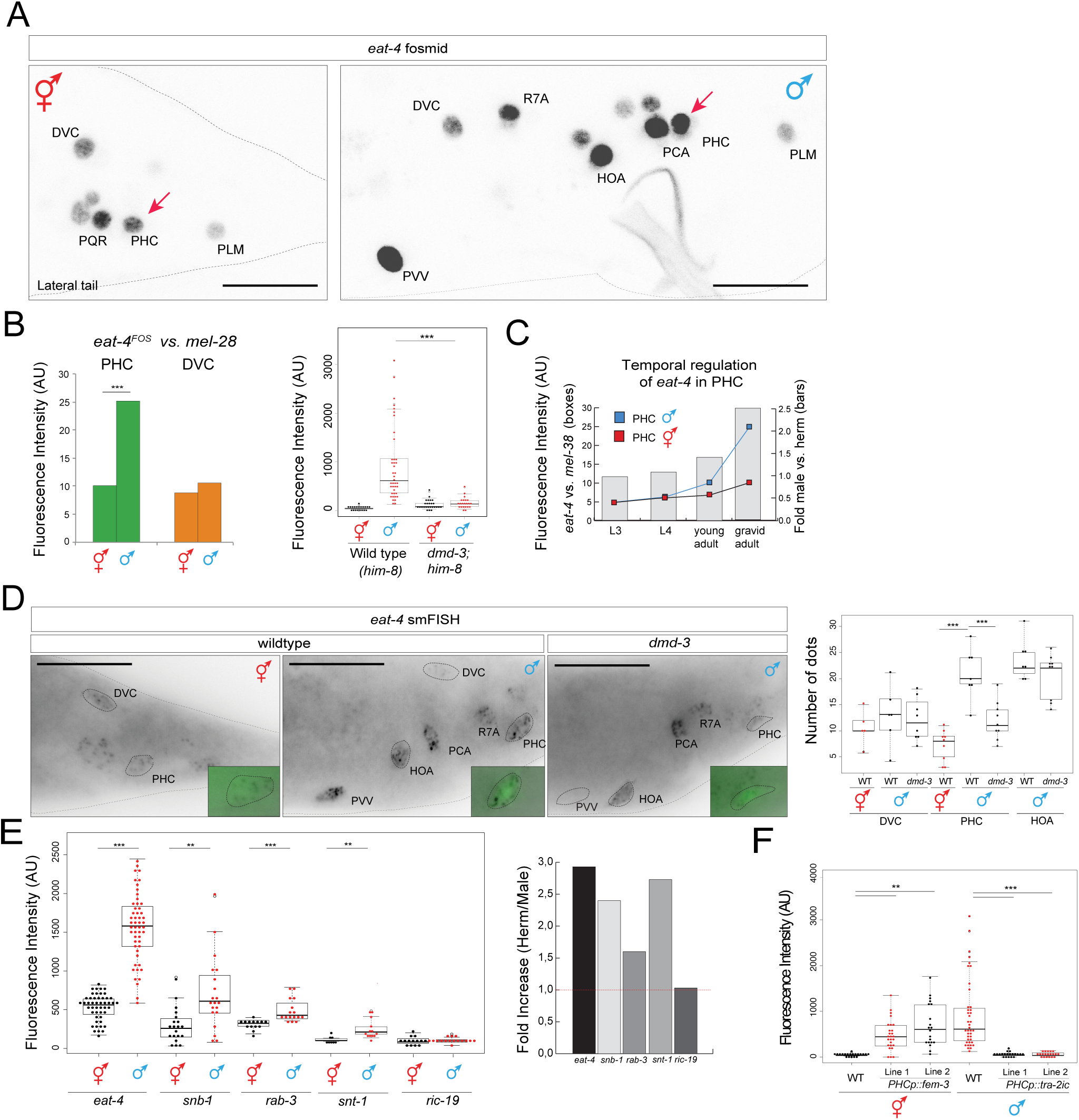
Scaling of synaptic vesicle components in PHC neurons. **A:** *eat-4/Vglut* fosmid reporter expression (*otIs518*) in the adult male tail. The complete set of all male-specific, *eat-4/Vglut-*expressing neurons will be published elsewhere. Scale bar: 20 µm. **B:** *eat-4/Vglut* fosmid (*otIs518*) reporter expression measured by absolute fluorescence (right panel) and normalized to expression of the *mel-28* nuclear envelope protein (*BN468*); *otIs520* reporter expression measured by absolute fluorescence in *him-8(e1489)* background and *dmd-3(tm2863);him-8(e1489)* (left panel). **C:** Temporal dynamics of scaling of *eat-4/Vglut* fosmid (*otIs518*) reporter expression. **D:** smFISH analysis of endogenous *eat-4/Vglut* expression in young adult animals. PHC was marked with *otIs520* (green). Scale bar: 20 µm. **E:** Scaling of transcription of other vesicular markers, as assessed by SL2-based fosmid reporter expression [30]. **F:** *eat-4/Vglut* scaling is controlled cell-autonomously as shown by masculinization and feminization experiments. *otIs520* (*eat-4prom11::gfp*) was crossed with *PHCp::fem-3* (lines *otEx6879* and *otEx6980*) and with *PHCp::tra-2ic (lines otEx6881 and oEx6882)* strains. PHCp (*eat-4prom11Δ11* driver). Wildtype data showing in this plot is same as in panel B (left panel), genotypes were scored in parallel. Significance was calculated using student t-test,\*\*\**P* < 0.001, \*\**P* < 0.01.

We examined whether the increase in *eat-4/Vglut* expression and synaptic output is accompanied by the upregulation of other synaptic proteins. Using fosmid-based reporter genes, we find that transcription of the vesicular proteins synaptotagmin (*snt-1*), synaptobrevin (*snb-1*) and Rab-3 (*rab-3*) is also upregulated (Fig.3E). *ric-19*, a gene encoding the ortholog of ICA69, involved in dense core vesicle mediated secretion, shows no upregulation (Fig.3E). Taken together, we define here a previously unrecognized transcriptional scaling phenomenon in which the generation of novel synaptic contact is mirrored by a neuron-type specific transcriptional upregulation of vesicular proteins.

We again used cell-specific sex reversal experiments to test whether the scaling of synaptic vesicle machinery requires the presence of the entire male-specific circuitry that PHC becomes wired into or whether it is controlled cell autonomously, like PHC axon extension. We find that masculinization of PHC via *fem-3*-mediated degradation of TRA-1 in otherwise hermaphroditic animals results in scaling of *eat-4/Vglut* expression (Fig.3F). Conversely, feminization of PHC in otherwise male animals using the intracellular domain of TRA-2, *tra-2ic*, expressed specifically in PHC, results in a failure to upregulate *eat-4/Vglut* expression (Fig.3F).

### Sexually dimorphic neuropeptide expression in PHC

Fast synaptic transmission machinery is not the only neurotransmission-related feature that becomes modulated in male PHC neurons. Through an expression analysis of candidate genes, we also identified a FMRFamide neuropeptide-encoding gene, *flp-11*, with dimorphic expression in PHC. *flp-11* codes for at least four different peptides and is phylogenetically conserved in nematodes [20]. In addition to non-dimorphic expression in a number of tail neurons, *flp-11* is expressed in PHC neurons exclusively in males, but not in hermaphrodites (Fig.4A). *flp-11* is not expressed in either sex in early larval stages (L2, L3), after the birth of PHC in the L1 stage and before sexual maturation, but it becomes activated specifically in the PHC neurons in L4 males (Fig.4B).

**Fig. 4:**
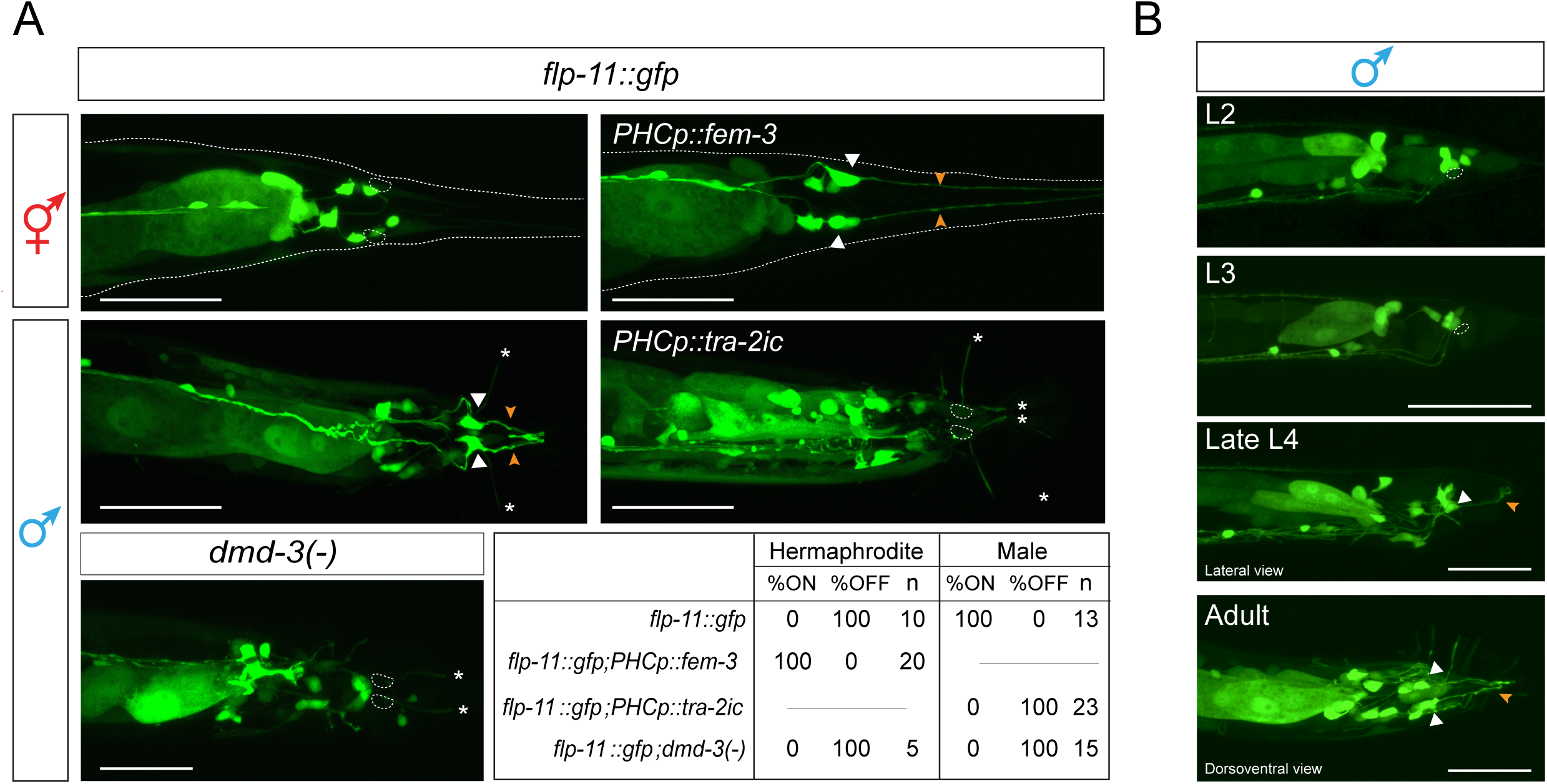
Sexually dimorphic neuropeptide expression in PHC. **A:** *flp-11* reporter expression (*ynIs40)* in wildtype hermaphrodites and males, in animals in which the sex of PHC was changed via *fem-3* or *tra-2ic* expression and in *dmd-3* mutants. The table indicates the percentage of neurons that show expression of the *flp-11* reporter in the different conditions assayed. N=number of animals. Scale bar: 50 µm. **B:** Temporal dynamics of *flp-11* reporter gene expression in larval and adult male stages. Cell bodies are labeled with white arrow, dendritic projections with orange arrow. White asterisks indicate ray projections. Scale bar: 50 µm.

Like PHC axon extension and scaling of synaptic vesicle machinery, the male-specific induction of *flp-11* expression is controlled cell-autonomously. Masculinization of PHC via *fem-3*-mediated degradation of TRA-1 in otherwise hermaphroditic animals results in induction of *flp-11* expression (Fig.4A) and, conversely, feminization of PHC in otherwise male animals using the intracellular domain of TRA-2, *tra-2ic*, expressed specifically in PHC, results in a failure to induce *flp-11* expression (Fig.4A).

*flp-11* mutants do not display defects in the vulva location behavior controlled by PHC (Fig.1C), suggesting either redundancy of *flp-11* function with other neuropeptides or that PHC may be involved, via FLP-11, in the regulation of as yet unknown male-specific behaviors.

### The *eat-4/Vglut* locus contains *cis-*regulatory modules that confer sexually dimorphic expression

As a starting point to dissect the mechanistic basis of regulation of PHC remodeling, we turned to the *eat-4/Vglut* locus and asked how its sex-specific scaling is controlled. We have previously dissected the *cis*-regulatory architecture of the entire *eat-4/Vglut* locus, defining modular elements that drive *eat-4/Vglut* expression, and hence glutamatergic identity, in distinct glutamatergic neuron types throughout the nervous system of the hermaphrodite [21]. A fragment of the *eat-4* locus (*“prom5”)* drives expression in the PHC neuron as well as multiple other glutamatergic neurons [21](Fig.5A). We dissected this *prom5* element to a 494 bp element (“*prom11*”) with remarkable sex-specificity. Not only does this element recapitulate the sex-specific scaling of the endogenous *eat-4/Vglut* locus in the PHC neurons, but it also shows sexually dimorphic expression in a single pair of head interneurons, the AIM neurons (Fig.5). Using the full length fosmid reporter construct, we have previously shown that *eat-4/Vglut* expression is constitutively expressed in AIM, but is downregulated specifically in the AIM neurons in males during sexual maturation (upon *eat-4/Vglut* downregulation, the AIM neurons become cholinergic instead; [22]). We find that the sex specificity of *eat-4/Vglut* expression is recapitulated by the *prom11* module not only in regard to scaling of *eat-4/Vglut* expression specifically in male PHC, but also in regard to the downregulation of *eat-4/Vglut* expression in the male AIM neurons (Fig.5C).

**Fig.5:**
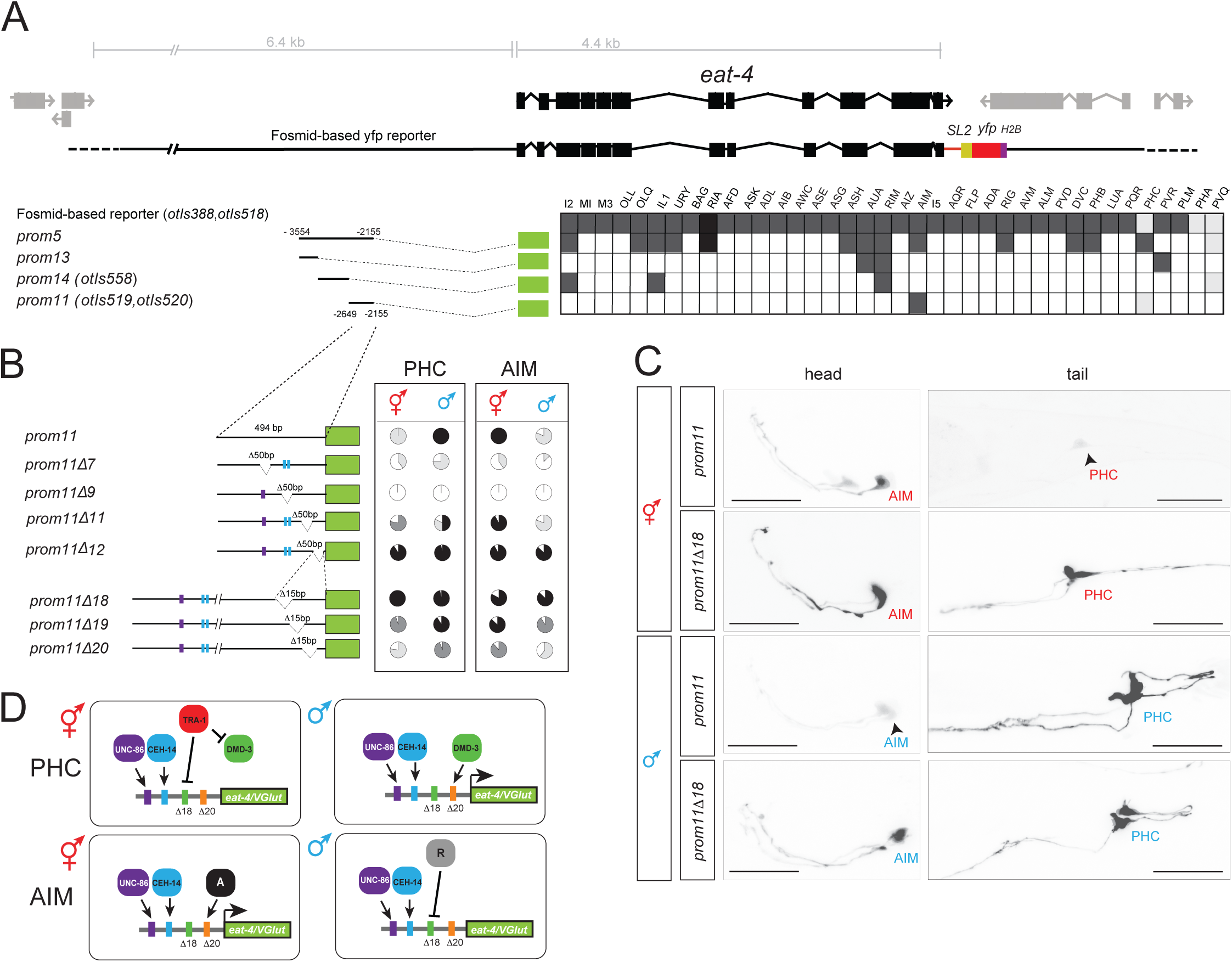
Analysis of the *cis*-regulatory control elements of the *eat-4* locus. **A:** Dissection of the *eat-4prom5* promoter. In the analysis of expression between two and three lines (n > 10) were scored for expression. The different shades of gray indicate the relative fluorescence intensity. Cell identifications were done crossing the lines with the *eat-4* fosmid reporter (*otIs518*). The asterisk indicates DIC identification. **B:** Deletion analysis of the *eat-4prom11* promoter. The pie charts indicate the percentage of neurons that express the array with the different shades of gray denoting the overall fluorescence intensity in AIM and PHC in both sexes. At least two lines were analyzed for each deletion construct. Color bars in the promoters refer to putative homeodomain binding sites: blue bar indicates the CEH-14 binding site according to the ModEncode consortium; purple bar indicates the putative UNC-86 binding site. **C:** *eat-4prom11* and *eat-4prom11Δ18* reporter expression in the head and tail of hermaphrodite and young adult males. Scale bar: 25 μm. **D:** Summary schematic of *eat-4/Vglut* regulatory logic. Note that the same elements are required to achieve activator and repressor effects in the PHC and AIM neurons, but with opposite sexual specificity. In the AIM neurons, unknown factors “A” (for activator) and “R” (for repressor) may be different DMD transcription factors. “A” could also be TRA-1, but now working, in this context, as an activator. “R” may be repressed by TRA-1 in hermaphrodite AIM.

How is the expression of the module controlled? We had previously shown that terminal differentiation of both the AIM and PHC neurons in hermaphrodites each require the LIM homeobox gene *ceh-14*, the *C. elegans* ortholog of vertebrate Lhx3/4 and its presumptive partner, the Brn3-like POU homeobox gene *unc-86* [21]. We examined whether *ceh-14* and *unc-86* not only control *eat-4/Vglut* expression in hermaphrodite PHC, as previously reported [21], but are also required for *eat-4/Vglut* expression (and its scaling) in males, and indeed find this to be the case (Suppl. Fig.S1). We identified predicted binding sites for UNC-86 and CEH-14 in the *eat-4prom11* element and found that mutations of each predicted binding site resulted in loss of expression of the *prom11* module in both AIM and PHC (Fig.5B). The authenticity of the CEH-14 binding site is corroborated by the ModEncode consortium, which mapped a CEH-14 binding peak to the location of the *prom11* element [23].

To decipher the *cis*-regulatory logic of sex-specific modulation of *eat-4/Vglut* expression in PHC (and AIM), we further dissected the *prom11* module by introducing deletions throughout this module. Our analysis was guided by two distinct hypotheses for how sex-specific regulation may be achieved: (1) sex-specific repressor elements may counteract the non-sex specific activity of *unc-86* and *ceh-14*; deletion of such repressor elements should result in depression of *eat-4prom11* expression in the opposite sex (i.e. derepression of *gfp* expression PHC of hermaphrodites). (2) sex-specific activator element(s) may assist *unc-86* and *ceh-14* to provide sex-specific upregulation of *eat-4/Vglut* expression; deletion of such elements should result in a failure to scale reporter gene expression. Through the introduction of deletions in the *prom11* module we found evidence for both mechanisms (Fig.5B). Aside from the presumptive UNC-86 and CEH-14 binding sites (*prom11Δ7* and *prom11Δ9*, respectively), we identified a small element of 15 bp that when deleted (in *prom11Δ18)*, results in derepression of *eat-4/Vglut* in the PHC neurons of hermaphrodites (**Fig.5B,C**). The hermaphrodite-specific TRA-1 protein mentioned above, commonly thought to be a repressor protein [24], could potentially acts through this *cis*-regulatory element (however, we could not find any canonical TRA-1 binding site contained in *prom11*). Notably, the *Δ18* deletion in *prom11* also results in derepression of reporter gene expression in the AIM head neuron, but with opposite sex-specificity. While the *prom11* construct is only expressed in AIM in hermaphrodites, the *Δ18* deletion results in derepression in males (Fig.5B,C).

Importantly, in addition to the negative regulatory element that prevents the *prom11* module to be expressed in hermaphrodite PHC, we also identified an element that is required, in addition to the CEH-14 and UNC-86 sites, for *eat-4/Vglut* scaling in PHC in males (Fig.5B,C; *Δ20*). Like the repressor element *Δ18* described above, the *Δ20* activator element also has an activating effect in AIM, but again with the opposite sexual specificity. The *Δ20* deletion results in a failure to activate reporter gene expression in hermaphrodite AIM (Fig.5B,C).

In conclusion, these findings suggest that sex-specificity of *eat-4/Vglut* scaling in PHC is achieved by combination of (a) non-sex specific activation, (b) specific repression in hermaphrodites (possibly via TRA-1) and (c) sex-specific activation in males. A similar activator/repressor logic appears to also act in the AIM neurons, but with opposite sexual specificity.

### *dmd-3* affects scaling of *eat-4/Vglut* expression and remodeling of all other dimorphic PHC features

To identify trans-acting factors that may control the male-specific upregulation of the *cis*-regulatory module of the *eat-4/Vglut* locus mentioned above, we turned to the phylogenetically conserved family of Doublesex/DMRT transcription factors, of which there are 11 homologs encoded in the *C. elegans* genome. Sexually dimorphic functions have been identified for five of these genes so far (*mab-3, mab-23, dmd-3, dmd-5, dmd-11*)[9, 10]. In all cases, these factors act in a male-specific manner to specify male-specific features and we therefore considered them as candidates for sex-specific scaling of *eat-4/Vglut* expression in male PHC neurons. We analyzed the impact of all *dmd* genes on *eat-4/Vglut* expression for which mutant alleles were available (data not shown). Of all genes tested, we found one gene with an effect on *eat-4/Vglut* expression: Loss of *dmd-3* results in a failure to scale *eat-4/Vglut* expression in PHC (Fig.3B,D). This effect matches the *Δ18* deletion within the *prom11* module, suggesting that DMD-3 may act directly or indirectly through this module.

We next tested whether the activity of *dmd-3* is restricted to scaling *eat-4/Vglut* expression, or whether *dmd-3* may be a “master regulator” of all PHC remodeling events that we described above. We find that in *dmd-3* mutants, the male-specific axon extension of PHC neurons fails to occur (Fig.2B, Fig.6B). Moreover, expression of the FMRFamides encoded by the *flp-11* locus does not become induced in PHC during sexual maturation of the male (Fig.4A). We conclude that *dmd-3* affects all currently measurable aspects of male-specific PHC differentiation.

**Fig.6:**
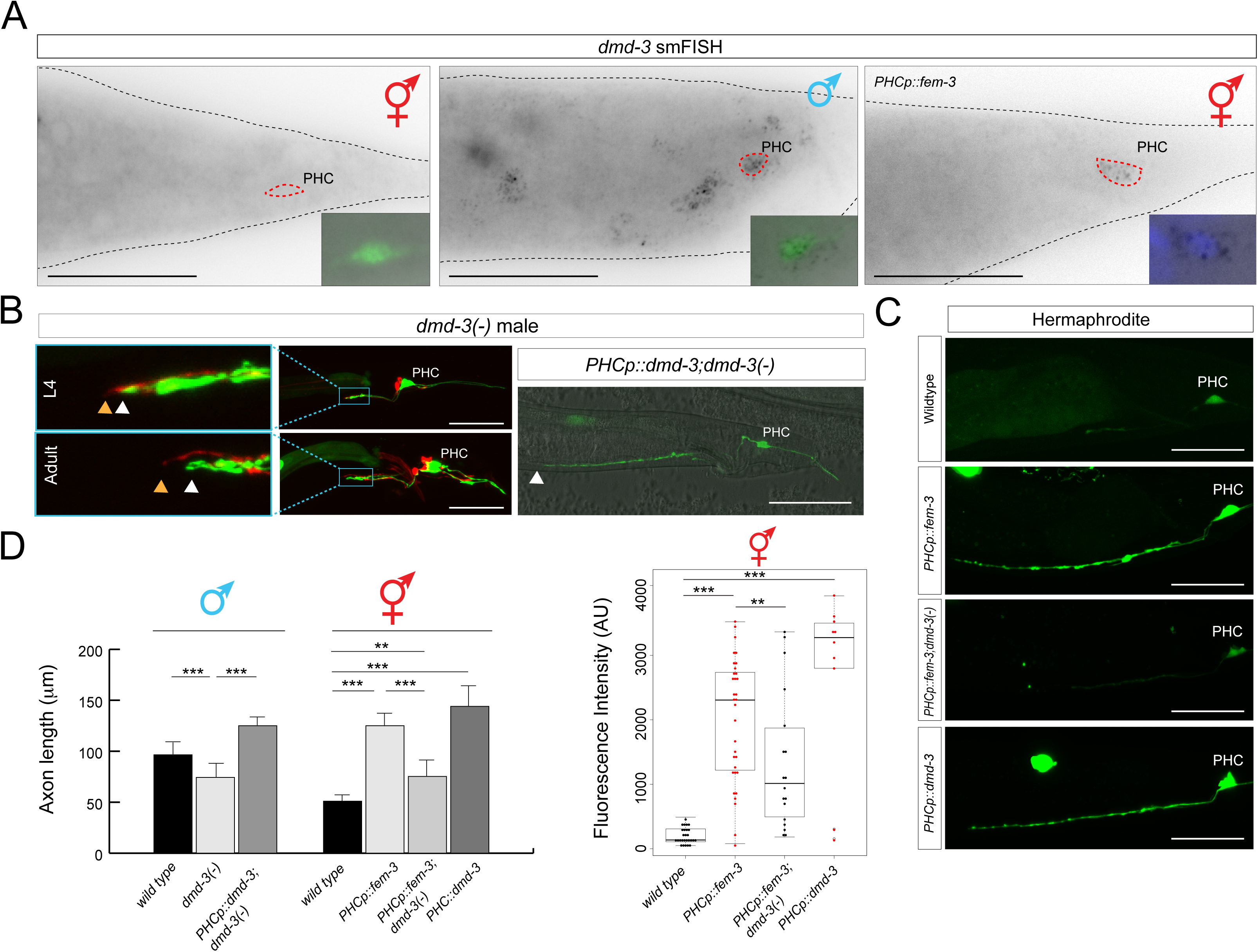
*dmd-3* expression and function. **A:** smFISH analysis of endogenous *dmd-3* expression in young adult animals. No expression is observed in hermaphrodites; in males, expression is observed in multiple cells including PHC, marked with the transgenic array *otIs520*. Masculinization of PHC (via *PHCp::fem-3*) in otherwise hermaphroditic animals activates *dmd-3* transcription. The inset shows the PHC nucleus stained with DAPI. More than 10 animals were scored for presence of dots in each condition and under each condition all animals showed the same staining patterns relative to one another. Scale bar: 50 µm. **B:** PHC axons fails to extend in *dmd-3(tm2863)* mutant males and these defects are rescued by PHC-specific expression of *dmd-3* [*oExt6908 (eat-4p11Δ11::dmd-3)*]. Scale bars: 50 µm. Wildtype and *PHCp::fem-3* data showing in this plot is the same as in **Fig.2B** for the adult stage. See panel D for quantification. **C:** Ectopic expression of *dmd-3* in the PHC neurons of hermaphrodites is sufficient to scale *eat-4/Vglut* expression (4^th^ panel) and *dmd-3* is required in hermaphrodites for the axon extension conferred by masculinization of PHC (2^nd^ and 3^rd^ panel). Transgenic array names: *otEx6879, otEx6880 (eat-4p11Δ11::fem-3); otEx6908(eat-4p11Δ11::dmd-3).* See panel D for quantification. **D:** Quantification of the axon extension and *eat-4/Vglut* scaling in the different conditions showed in previous panels. Significance was calculated using student t-test, \*\*\**P* < 0.005, \*\**P*< 0.05.

### *dmd-3* acts cell-autonomously in PHC and is sufficient to induce male-specific PHC differentiation

Upon sexual maturation, the entire male tail undergoes remodeling to form a copulatory structure [25]. The most posterior hypodermal cells in the tail define a specialized, sexually dimorphic structure in which cells fuse and retract in the male, changing their shape from a tapered cone to a blunt dome [25]. *dmd-3* has previously been shown to be expressed in hypodermal cells and required in the hypodermis for tail tissue remodeling [26, 27]. To examine whether *dmd-3* acts in the hypodermis or in the PHC neurons, we first examined the neuronal expression pattern of *dmd-3.* Using single molecule *in situ* hybridization, we find that *dmd-3* is expressed in a sex-specific manner not only in the male hypodermis, as previously reported[26, 27], but also in the PHC neurons. *dmd-3* expression in PHC is only observed in males, but not hermaphrodites (Fig.6A). Male-specificity is controlled cell-autonomously by *tra-1*, because masculinization of PHC using *fem-3* expression results in the induction of *dmd-3* expression in the PHC of hermaphrodites (Fig.6A). This derepression is functionally relevant, since both the axon extension and *eat-4/Vglut* scaling observed in transgenic hermaphrodites in which we masculinized PHC via *fem-3* expression requires *dmd-3* (Fig.6C,D).

To further corroborate that *dmd-3* indeed acts in PHC, we generated transgenic animals in which we expressed *dmd-3* under control of the PHC (and AIM)-specific fragment of the *eat-4/Vglut* locus. This construct rescues the axon extension defect of the PHC neurons in *dmd-3* mutant males (Fig.6B) as well as the *eat-4/Vglut* scaling defects of *dmd-3* mutants (Fig.6D).

Since the driver used to express *dmd-3* is not sex-specific (Fig.5D), we could also assess whether ectopic expression of *dmd-3* in hermaphrodites, achieved by this transgene, is sufficient for axon extension of PHC and scaling of *eat-4/Vglut* expression in a hermaphrodite. We indeed find that *dmd-3* is sufficient to induce these PHC features in hermaphrodites (Fig.6C,D). We conclude that *dmd-3* is both required and sufficient to autonomously specify the male-specific identity differentiation program of PHC and that its sex-specificity is controlled by sex-specific repression via TRA-1.

## DISCUSSION

Sexually dimorphic neuronal projection patterns have been observed in several distinct nervous systems from flies to human [2, 3, 28]. However, the simplicity and well-described nature of the *C. elegans* nervous system enables studies of neuronal sexual dimorphism with unrivaled anatomical, cellular and molecular resolution. The case of the PHC neurons particularly stands out because among the sex-shared neurons, PHC displays the largest extent of sexually dimorphic connectivity [6]. Other sensory neurons, like PHA or PHB also switch synaptic partners between the different sexes, but they do not differentiate into as densely connected hub neurons. One remarkable aspect of this sex-specific differentiation process is its cellular autonomy and hence independence from other network components. This autonomy extends to the newly discovered transcriptional scaling phenomenon of synaptic vesicle machinery that accompanies the synaptic remodeling of the PHC neurons. Cellular differentiation is often accompanied by the scaling of specific subcellular structures and their molecular components in order to adapt to the specialized needs of a cell [29], but such scaling phenomena have not previously been described in the context of the synaptic vesicle machinery. To the contrary, the expression levels of panneuronal genes, such as synaptic vesicle components, have been shown to be particularly resilient to genetic perturbations, a robustness that is ensured by multiple redundant *cis*-regulatory elements present in panneuronal gene loci [30].

We have shown here that a single transcription factor, *dmd-3,* is required and alone sufficient to trigger the male-specific program of this neuron, including the scaling of the synaptic machinery. The cellular context dependency of *dmd-3* function is striking. As previously shown, *dmd-3* is required in skin cells for their remodeling during male tail morphogenesis [26, 27], in male-specific cholinergic ray neurons for their differentiation [31] and, as we have shown here, is independently required in a sex-shared neuron to remodel specific anatomical and molecular features. The specificity of action of *dmd-3* is determined by cell type-specific combinations of cofactors. In the case of the hypodermis and male-specific cholinergic neurons, DMD-3 interacts with another DMD factor (MAB-23)[26, 31], which we find to have no impact on PHC differentiation (data not shown) while in the PHC neurons *dmd-3* cooperates with the *unc-86* and *ceh-14* homeobox genes. For *dmd-3* to exert its effect on *eat-4/Vglut* scaling, *unc-86* and *ceh-14* are required as permissive co-factors. One unanticipated conclusion from the *cis*-regulatory analysis of the synaptic scaling process is that the sex-specificity of this transcriptional regulatory phenomenon is not merely assured by one sex-specific regulatory factor that operates in one sex to either activate or repress a specific gene. Rather, sex-specific *eat-4/Vglut* expression requires sex-specific repression in one sex (mediated by a specific *cis*-regulatory element that may be directly or indirectly controlled by TRA-1) and sex-specific activation in the opposite sex (mediated by a specific *cis*-regulatory element that may be directly or indirectly controlled by DMD-3).

We anticipate that similar intersectional regulatory scenarios apply in other nervous systems where broadly acting, sex-specific regulatory factors, such as sex hormone-activated nuclear hormone receptors intersect with cell-specific regulatory factors to trigger cell-specific responses to sex hormones.

## EXPERIMENTAL PROCEDURES

### Strains and transgenes

Wild type is strain *him-5(e1490)V* or *him-8(e1489)IV* when indicated. Strains were maintained by standard methods. Mutant strains used in this study: *flp-11(tm2706)V*, *dmd-3(tm2863)V*, *mab-3(a1240)II*, *mab-23(gk664)V*, *dmd-5(gk408945)II, dmd-6(gk287)IV, dmd-7(ok2276)V, dmd-8(tm5083)V, dmd-9(tm4583)IV, dmd-10(gk1131)V, dmd-11(gk552)V, eat-4(ky5)III.* A list of transgenes can be found in **Suppl. Table S1**.

### Male mating assay

Mating assays were done as described previously [9, 13, 32](Liu and Sternberg 1995, Garcia et al 2007 and Oren-Suissa 2016). Early L4-stage males were transferred to a fresh plates and kept apart from hermaphrodites until they reach sexual maturation (24 hours). Single virgin males were assayed for their mating behavior in the presence of 10-15 adult *unc-31(e928)* hermaphrodites on a plate covered with a thin fresh OP50 lawn. *unc-31* hermaphrodites move very little, allowing for an easy recording of male behavior. Mating behavior was scored within a 15 min time window or until the male ejaculated, whichever occurred first. Males were tested for their ability to locate vulva in a mating assay, calculated as location efficiency (L.E.)[33]. The number of passes or hesitations at the vulva until the male first stops at the vulva were counted. Location Efficiency = 1 / # encounters to stop. *PHCp::HisCl1* transgenic animals were transferred to NGM plates containing 10mM histamine a day prior to the mating assay [9, 11].

### Harsh touch assay

Harsh touch assays were done as previously described [34]. L4 animals were separated by sex and transferred to either NGM plates or NGM plates containing 10mM histamine (both seeded with OP50) and allowed to mature to adulthood (24 hours). For the assay, adults were transferred to NGM or NGM plus histamine plates seeded with a thin fresh OP50 lawn. Harsh touch stimulus was administered to the anus of the worm using a platinum wire pick while the worm was stationary. An animal that initiated abrupt forward locomotion in response to harsh touch was considered “responding.” Each animal was only assayed one time.

### DNA constructs

The *eat-4prom11* was generated by PCR and cloned into pPD95.75. It contains the genomic region between -2680 and -2155bp from ATG of the *eat-4/Vglut* locus. The *eat-4prom11* deletions were generated by mutagenesis with the QuickChange Site-Directed Mutagenesis kit (Stratagene). The plasmids were injected in *him-5(e1490)* at 50 ng/μl, using *unc-122::gfp* (50 ng/μl) as a co-injection marker.

To generate the *eat-4p11Δ11::fem-3* and *eat-4prom11Δ11::tra-2ic*, the *eat-4prom11Δ11* fragment was cloned into the plasmids *pMO12 (inx-18p::fem-3::sl2::tagRFP)* and *pMO32 (inx-18p::tra-2ic::sl2::tagRFP)*[9] by Restriction Free-cloning by substituting the *inx-18* promoter. The DNA was injected into *him-5(e1490)* at 20 ng/μl, using *unc-122::gfp* (50 ng/μl) as a co-injection marker.

To generate the *eat-4p11Δ11::dmd-3* the cDNA of *dmd-3* was amplified from total RNA extracted from *him-5(e1490)* animals using the SuperScript III One-Step RT-PCR system (Invitrogen^™^) with the primers: 5’-ATGAATATAGAGGAAATTC-3’ and 5’-TTACTCATTTTTATGGGGAC-3’. The *dmd-3* cDNA was subcloned into *eat-4prom11Δ11::fem-3::sl2::tagRFP*, replacing the *fem-3* cDNA by RF-cloning. The DNA was injected into *otIs520*;*him-5(e1490)* at 20 ng/μl, using *unc-122::gfp* (50 ng/μl) as a co-injection marker.

The *eat-4prom11Δ11::HisCl::sl2::gfp* (*PHCp::HisCl*) was generated by subcloning the *eat-4prom11Δ11* fragment into the plasmid *pNP424* [9] digested with the restriction enzymes SphI and XmaI. The DNA was injected into *him-5(e1490)* at 50 ng/ul, using *unc-122::gfp* (50 ng/μl) as a co-injection marker.

### Single molecule FISH

smFISH was done as previously described [35]. Young adult animals were incubated over night at 37°C during the hybridization step. *dmd-3* and *eat-4* probes were designed by using the Stellaris RNA FISH probe designer and were obtained already conjugated with Quasar 670 and purified, from Biosearch Technologies. Both probes were used at a final concentration of 0.25 µM.

### Microscopy

Worms were anesthetized using 100mM of sodium azide (NaN_3_) and mounted on 5% agarose on glass slides. All images were acquired using a Zeiss confocal microscope (LSM880). Acquisition of several z-stack images was performed with the Micro-Manager software (Version 3.1). Image reconstruction was performed using ZEN software tool.

For the quantification fluorescence intensity of the *eat-4/Vglut* different arrays, a stack of images was acquired using a Zeiss confocal microscope (LSM880). The acquisition parameters were maintained constant among all samples (same pixel size and laser intensity). The fluorescence intensity mean was obtained with the ZEN software tool.

### DiD staining

Animals were collected and washed with M9 two times and then incubated for 45 minutes in DiD (1/2000) (Invitrogen^™^) at room temperature. Then, the DiD was washed twice with M9 and the animals were plate in regular NGM plates. For the imaging of adult males, L4 larvae were treated the day before. All the other stages were imaged 4 hours after treatment.

## ACKNOWLEDGEMENTS

We thank Qi Chen for generating transgenic strains and member of the Hobert lab for comments on the manuscript. Strains were provided by the CGC. This work was supported by the NIH (2R37NS039996) and the HHMI. M.O. received postdoctoral fellowship support from the EMBL and the HFSPO. E.A.B. received predoctoral fellowship support from the NIH (1F31NS096863-01).

## FIGURE LEGENDS

**Suppl.Fig.S1:**
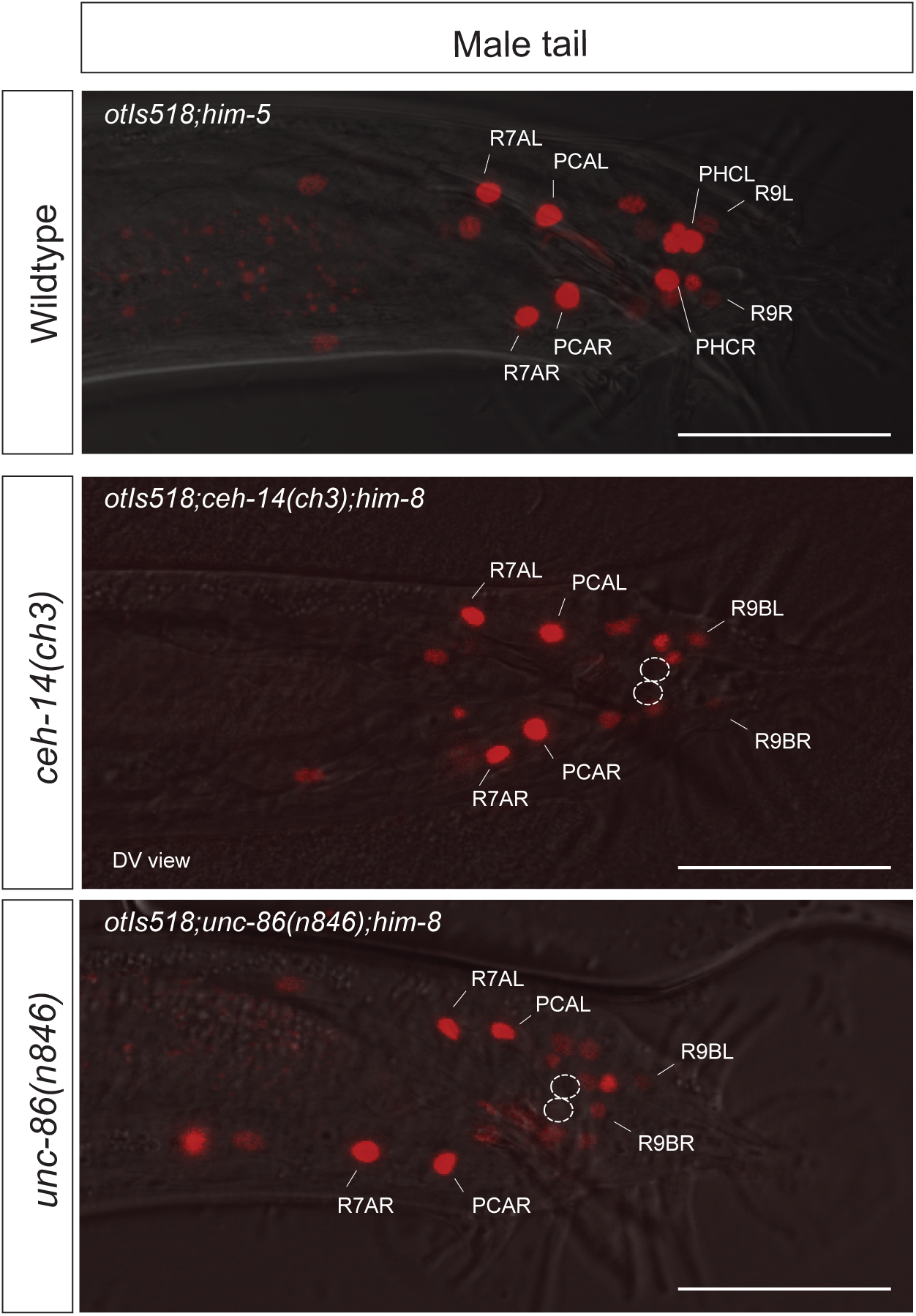
Effect of *ceh-14* and *unc-86* on *eat-4* scaling in PHC. *eat-4/Vglut* fosmid reporter expression (*otIs518*) in the adult male tail in wildtype and in *ceh-14(ch3)* and *unc-86(n846)* mutant backgrounds. 100% of animals show expression in PHC in wildtype animals (n>100); *ceh-14* and *unc-86* affect the expression of *eat-4* fosmid in PHC in 100% of the animals (n=5 for each genotype). Scale bar: 50 µm.

